# Robust TCR production for the structural study of TCR-pMHC complexes

**DOI:** 10.1101/2025.05.12.653033

**Authors:** Leire Oyon-Olea, María Gilda Dichiara-Rodríguez, Sandra Hervás, Elena Erausquin, Jacinto López-Sagaseta

## Abstract

A precise comprehension of how T cell receptors (TCRs) engage their antigens is pivotal for advancing basic research and T cell immunotherapy in cancer. While TCR refolding from inclusion bodies has greatly facilitated X-ray studies over the past decades, the procedures remain labor-intensive and can yield poorly. We have developed a simplified strategy for efficient production of soluble TCRs in CHO cells which, coupled with the removal of N-glycosylation, enable structural studies of TCR-pMHC complexes. An equivalent of just 20 ml of cell culture delivered sufficient deglycosylated TCR (dgTCR) to screen, upon complexation with a cognate pMHC, over 350 crystallization conditions and obtain a high-resolution dataset. This approach illustrates an effective alternative for TCR production to support studies devoted to research and development of TCRs.

## Introduction

T lymphocytes are key cellular components of the adaptive immune system, allowing responses against a vast array of antigens, including peptides derived from pathogens^1,2^ or tumors^3,4^, and even self-antigens^5–7^, and conferring immunological memory. More precisely, T cells recognize through their T cell receptors (TCRs) antigens when these are displayed on the surface of the body’s own cells, delivered by Major Histocompatibility Complex (MHC) molecules^8^.

Since recognition of peptide:MHC (pMHC) complexes by TCRs is crucial for proper immune function and health, understanding how these interactions occur provides fundamental insights into antigen-receptor specificity, T cell activation and immune memory formation. Therefore, this knowledge aids in the development of therapeutic strategies targeting infectious disease, cancer, autoimmune disorders, immunosuppression and vaccine development.

High-resolution structural characterization of TCRs and TCR-bound pMHC (TCR-pMHC) complexes requires high-quality soluble TCRs, which can be challenging to obtain. Various expression systems and strategies are used to produce TCR samples for crystallization studies, yet there still exist disadvantages that prevent high production yields or crystallization rates^9^.

*Escherichia coli*-based expression systems remain the most frequently used recombinant protein production organisms when protein crystallization and structural characterization is intended. However, the extraordinarily reducing environment of the bacterial cytoplasm drastically impairs the formation of disulfide bonds, which are essential for the proper folding of proteins like TCRs. Consequently, TCRs expressed in *E. coli* tend to form insoluble aggregates called inclusion bodies, resulting in low yields of correctly folded protein ^10,11^. Although biologically active proteins can be recovered from inclusion bodies through refolding protocols^12^, this process is challenging and labor-intensive, and may not yield sufficient protein for high-throughput crystallization screening.

Alternative strategies for soluble TCR production have been explored to avoid refolding after expression in *E. coli*. These include single-chain TCR versions ^13^, and stabilized TCR heterodimer constructs using jun/fos coiled-coil domains ^14^, a method which despite yielding soluble TCRs which have been proven to retain specific ligand binding activity, were difficult to crystallize.

Eukaryotic cells, such as mammalian or insect cells, can produce biologically active, properly folded recombinant proteins, including TCRs ^15,16^.However, intracellular folding processes in these cells can lead to post-translational modifications (PTMs), such as phosphorylation, acetylation or glycosylation. N-glycosylation usually occurs during protein translation at the Asn-X-Ser/Thr consensus sequence (where X is any amino acid except proline) and results in the addition of complex glycans. These glycans can be structurally heterogeneous and conformationally flexible, which can interfere with protein packing during crystal nucleation and hinder crystallization and structural studies. Enzymatic removal of glycans is commonly used to address this issue, but it can result in partially and heterogeneously deglycosylated protein samples.

To address these challenges and obtain high quality, and high-yield soluble TCR samples for crystallography, we have designed a straightforward strategy using mammalian CHO cells that greatly simplifies the end-to-end TCR production pipeline (Figure 1).

**Figure 1.**
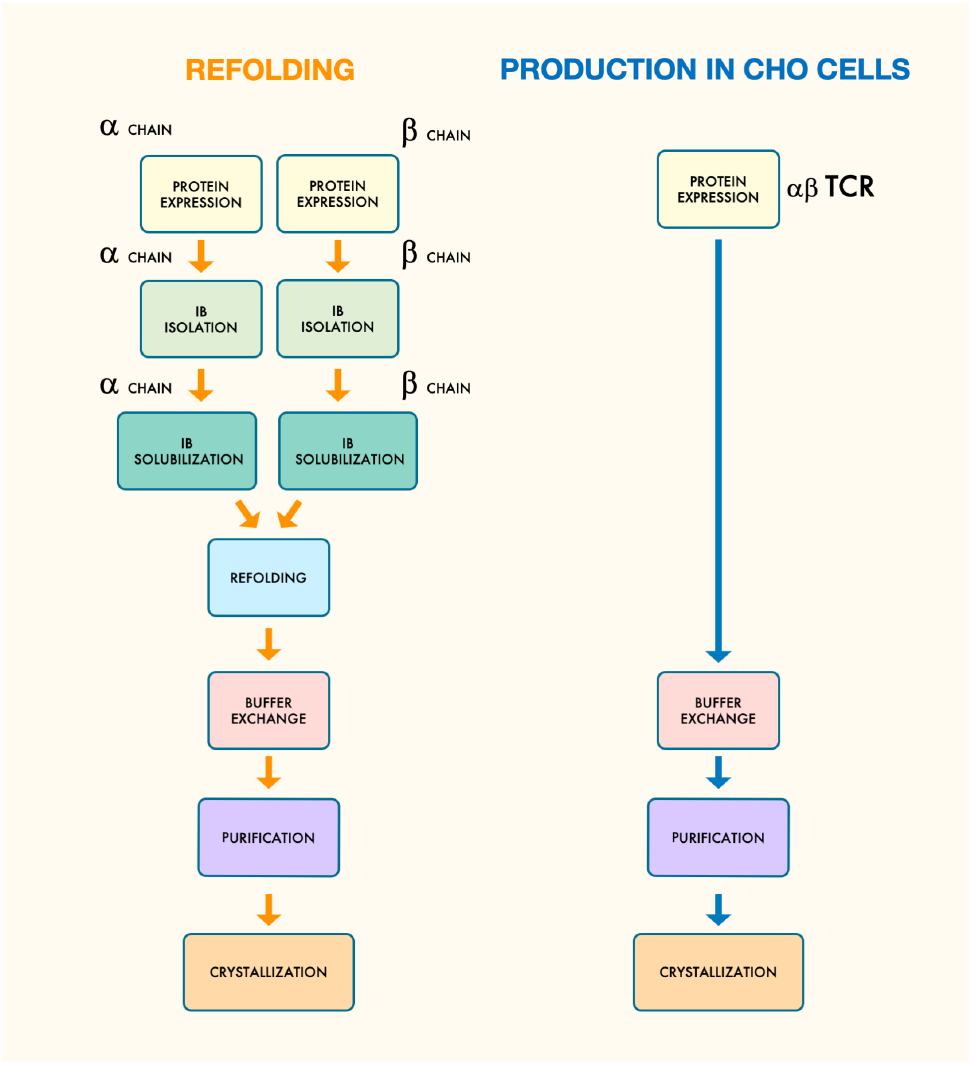
TCR production in CHO cells greatly simplifies the production and purification pipeline. The figure compares two methods for producing αβ T cell receptors (TCRs). Refolding approach (left side) involves a multi-step process and handling large volumes: α and β chains are expressed separately in bacteria. Proteins are recovered from inclusion bodies, solubilized, and refolded in vitro. This is followed by buffer exchange, which implies handling large volumes for dialysis, purification, and crystallization. CHO cell production (right side): The full αβ TCR is expressed directly in CHO cells, bypassing inclusion body (IB) isolation, solubilization, and refolding, and reducing significantly the volume to manipulate throughout the whole process, thereby streamlining the production of functional soluble TCR.

In addition, and importantly, to favor crystallization, our procedure completely prevents N-glycosylation at the TCR constant regions. In the work described herein, we have genetically engineered human TRAC*01 and TRBC2*01 sequences to remove N-glycosylation susceptible sites. More precisely, we propose the substitution of the Asn residues at these Asn-X (except Pro)-Thr/Ser sites for Gln residues (N28Q, N62Q and N73Q at the TCR alpha chain constant region, and N63Q at the TCR beta chain constant region) for high-yield native and soluble TCR protein production and their high-resolution structural characterization.

## Results

### Deglycosylated TCR is efficiently produced in CHO-S cells

As part of ongoing studies on the development of a TCR with tumor suppression properties, we initiated structural studies to define the structural basis for their ability to recognize an immunodominant tumor antigen peptide presented by HLA-A*02:01. Previous attempts using BL21(DE3) cells and codon-optimized genes to generate inclusion bodies for subsequent refolding procedures led to the absence of expression of the TCR alpha chain. Therefore, we explored the use of a CHO cell line as eukaryotic expression system.

Figure 2a summarizes the design of the TCR alpha and beta chain constructs, including N28Q/N62Q/N73Q and N63Q substitutions in the human TRAC and TRBC sequences, respectively, to prevent N-glycosylation of the TCR constant regions.

**Figure 2.**
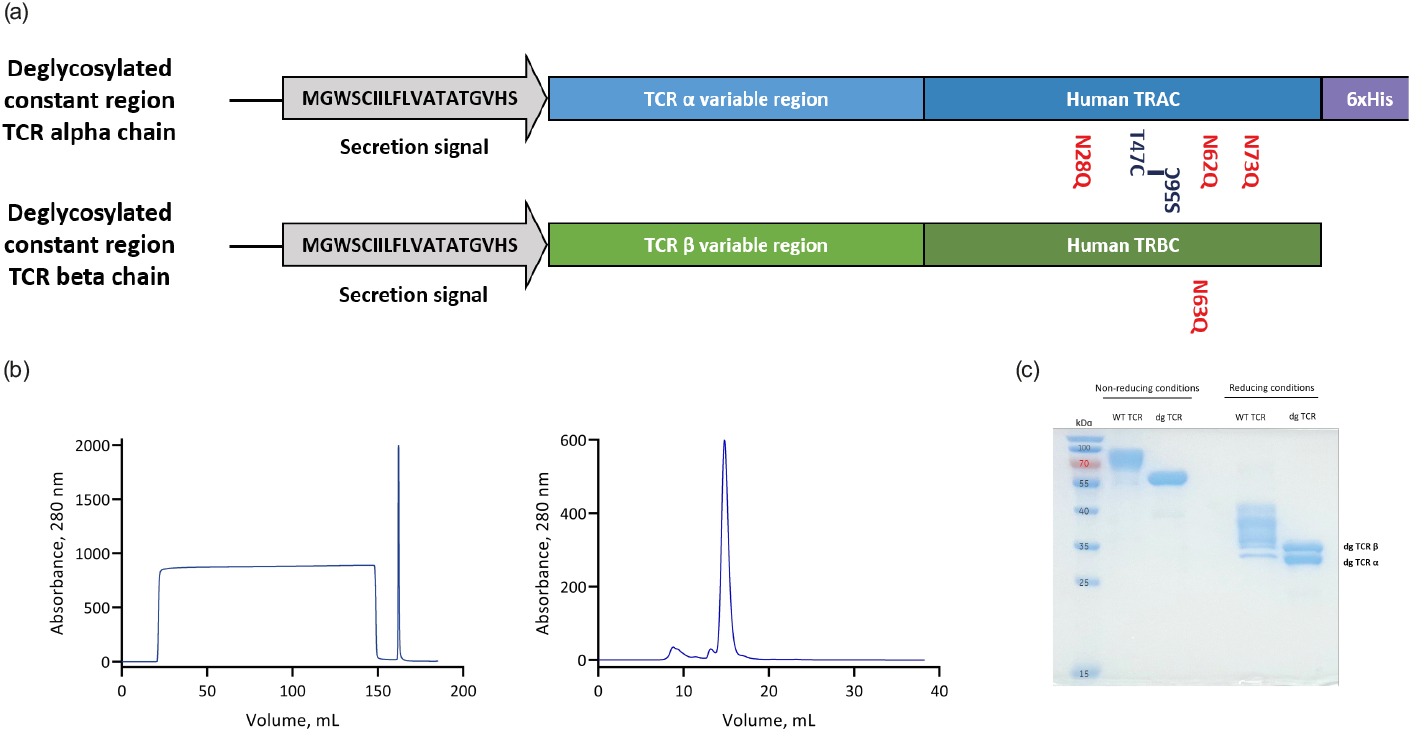
Production of *wild type* and dgTCR. (a) Schematic representation of the constructs cloned into the pcDNA3.4 vector for each TCR *wild-type* α/β and dgTCR α/β chain. The artificial substitutions introduced into the human TRAC and TRBC sequences are shown in red. (b) IMAC chromatography of the dgTCR purification using a HisTrap cartridge (left), and size exclusion chromatography of the HisTrap-purified dgTCR (center). (c) SDS-PAGE comparing *wild-type* TCR samples vs dgTCR eluted from the SEC chromatography shown in (b). Samples were run both in non-reducing (left) and reducing conditions to break interchain disulfide bridges (right).

CHO-S cells were co-transfected with the deglycosylated TCR (dgTCR) alpha and beta chains (1:1 ratio) to induce transient expression of the respective heterodimers. Eight days after transfection, the secreted αβ heterodimers were purified from the culture supernatants.

The clarified supernatant was dialysed against HBS pH 7.4 for 48 hours prior affinity chromatography using HisTrap cartridges on an ÄKTA start system (Figure b). His-tagged heterodimers were dissociated from the column using HBS pH 7.4, 200 mM imidazole, and the eluted protein was directly loaded onto a Superdex 200 10/300 GL column on an ÄKTA pure system to remove further impurities and buffer exchange the sample into TBS pH 7.4 (Figures 2b, c).

### dgTCR yields high crystallization rates

To generate the trimolecular dgTCR:pMHC complexes, pure dgTCR samples were combined with equimolar amounts of pMHC (HLA-A*02:01) at room temperature to allow complex formation.

In our first attempts with the wild type TCR counterpart, only one hit was found at day 14^th^, with an appearance of a profusion of micro crystals, out of 192 conditions (ProPlex HT-96 and PACT *premier* HT-96). Given this poor outcome, we sought to enhance the crystallization rate by preventing the heterogeneous and branched N-glycosylation.

Therefore, we substituted Asn residues with Gln residues at the N-glycosylation sites in the TCR constant regions. This approach was well-tolerated by the CHO cells, obtaining a yield of 26.4 mg/liter of pure dgTCR.

Starting from this material, 0.5 mg of pure dgTCR were taken apart for complexation with the pMHC. The remaining purified protein (∼ 2.1 mg) was aliquoted and stored frozen. Subsequent crystallization trials with the dgTCR-pMHC complex produced 17 and 11 hits with ProPlex HT-96 and PACT *premier* HT-96 kits, respectively. Two additional screens (SG1 Screen HT-96 and JCSG-*plus*) yielded 16 and 8 hits, respectively. Therefore, a total of 52 initial crystallization hits were obtained with the dgTCR:pMHC complex, which represents a crystallization rate of 13.5% (Figure 3). Some crystals were optimized by varying precipitant concentration and/or buffer pH, producing large or medium sized, single, diffraction-ready crystals (Figure 4a). Similarly, a related TCR with a different CDR3 alpha sequence yield 15, 46, 19 and 10 hits, respectively, in the ProPlex HT-96, PACT *premier*, SG1 Screen HT-96 and JCSG-*plus* screens, respectively, corresponding to a 23.4% of crystallization rate. These results confirm and strengthen the high crystallizability of the dgTCR.

**Figure 3.**
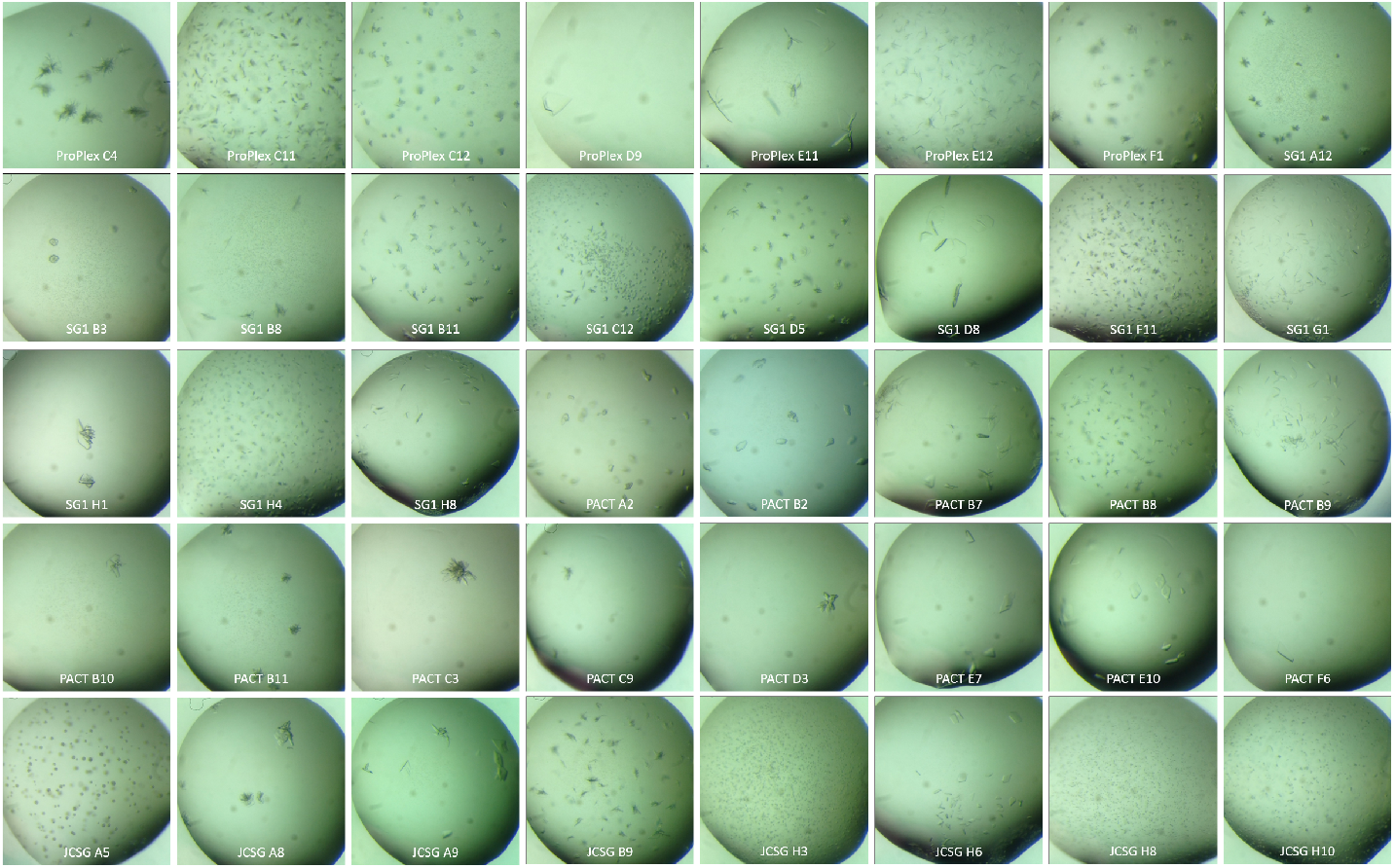
dgTCR:pMHC complex formation and crystallization. Examples of the crystals obtained from various crystallization screens with the dgTCR:pMHC complex. Each panel represents a different condition from various crystallization screens (ProPlex, SG1, PACT, JCSG). Images were taken 24 hours after setting up the crystallization screen.

**Figure 4.**
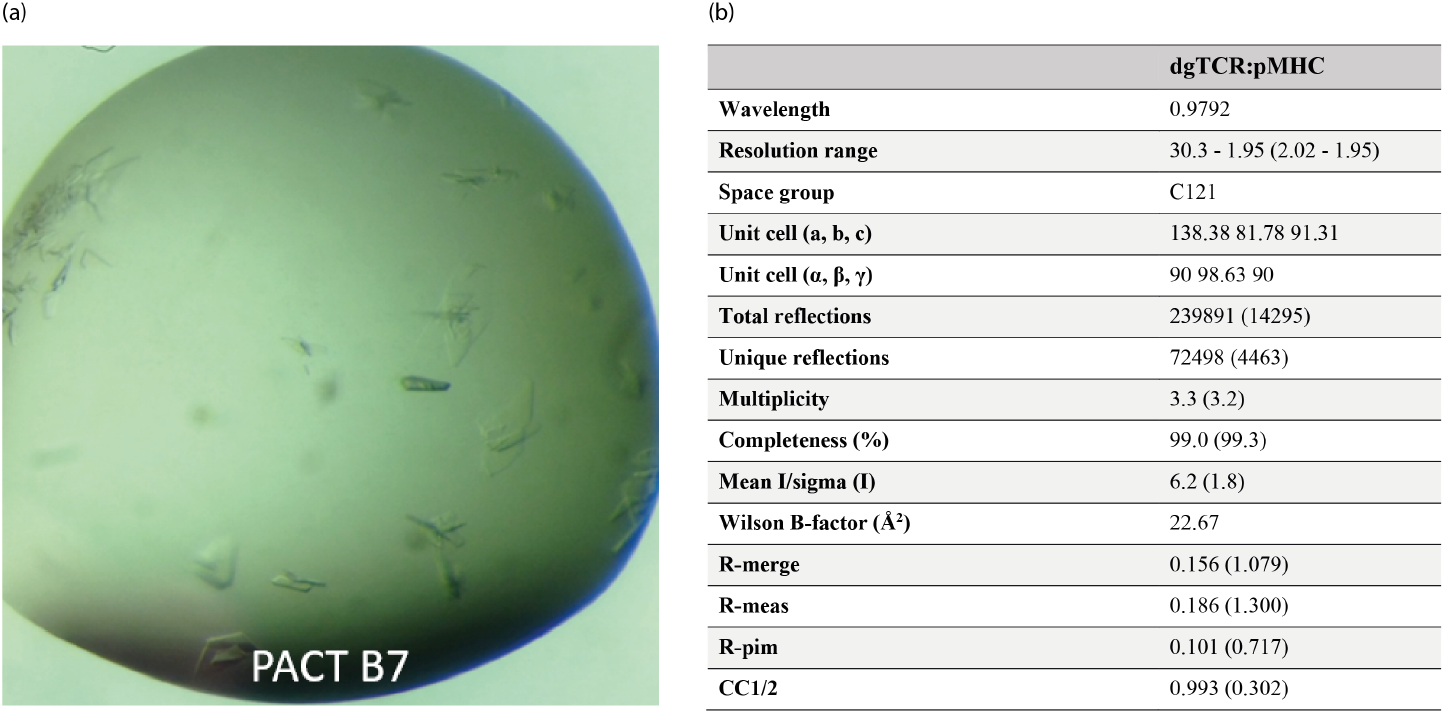
Diffraction-grade dgTCR:pMHC complex crystals. (a) Close-up view of the crystals grown in condition PACT B7 (0.2M NaCl, 0.1M MES pH 6.0, 20% w/v PEG 6000) as part of the original crystallization screening. Several plate-like crystals are visible within the drop (b) Diffraction data statistics of a single and native dataset obtained with a single crystal from the drop shown in the picture. Values in parentheses indicate data in highest-resolution shell.

### dgTCR produces high resolution datasets for structure determination

Irregular or rhomboid-shaped thin crystals of dgTCR:pMHC grew in 0.2 M sodium chloride, 0.1 M MES pH 6.0, 20% w/v PEG 6000. Crystals were harvested, soaked in their crystallization medium supplemented with 20% glycerol and cryoperserved in liquid nitrogen prior to diffraction analyses.

X-ray diffraction of dgTCR:pMHC crystals resulted in the acquisition of several datasets which allowed ternary complex structure solution with a resolution of 1.95 Å (Figure 5b). The dgTCR:pMHC complex crystallized in space group C121 with a single TCR:peptide/MHC-I molecule per asymmetric unit. The structure was solved via molecular replacement. Structure refinement combined with model building led to the final electron density maps and atomic coordinates of the dgTCR:pMHC complex (Figure 5a).

**Figure 5.**
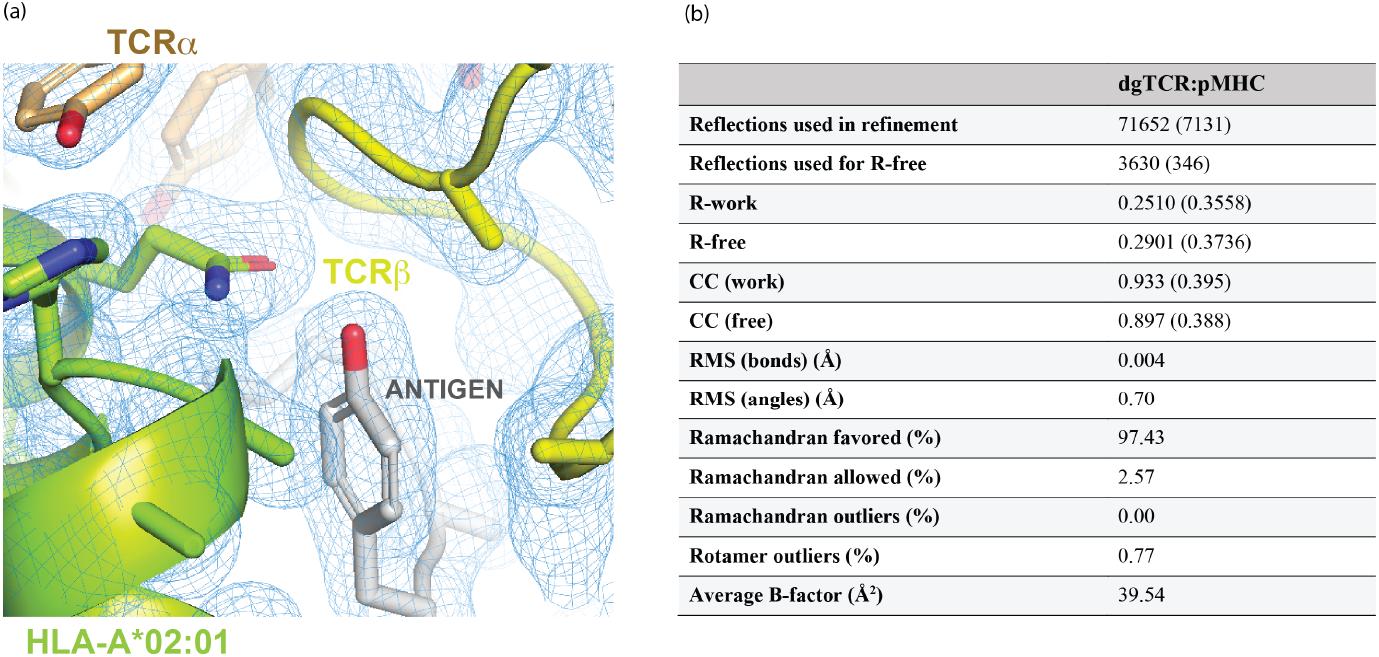
Structure resolution of the deglycosylated TCR:pMHC complex. (a) Close-up view of the dgTCR complexed with the antigen peptide presented by HLA-A*02:01. The blue mesh represents the electron density map (2Fo–Fc), confirming the fit of the structure to the experimental data. The TCR α (orange) and β (yellow) chains bind on top of the antigen (grey)-MHC (green) scaffold, following an overall canonical binding pattern. (b) Crystallographic refinement statistics table for the dgTCR:pMHC complex. Values in parentheses indicate data in highest-resolution shell.

Recognition of pMHC-I by the dgTCR follows an overall canonical TCR:peptide/MHC-I docking pattern, where the dgTCR binds, through its CDRs, on top of the peptide-binding scaffold formed by the MHC-I molecule. Therefore, the biological significance of the resolved structure confirms that deglycosylation of the TCR constant regions does not alter TCR function and enables proper docking on top of the MHC molecule as well as adequate stacking of the complex for crystallization. In addition, compared to other deposited TCR:pMHC structures, our deglycosylated structure diffracted at a higher resolution than the average, demonstrating its suitability for X-ray diffraction analyses (Figure 6a). A similar trend is observed when resolution of our dgTCR:pMHC structure is compared with other deposited TCR complexes bound to the same MHC allele (HLA-A*02:01, Figure 6b).

**Figure 6.**
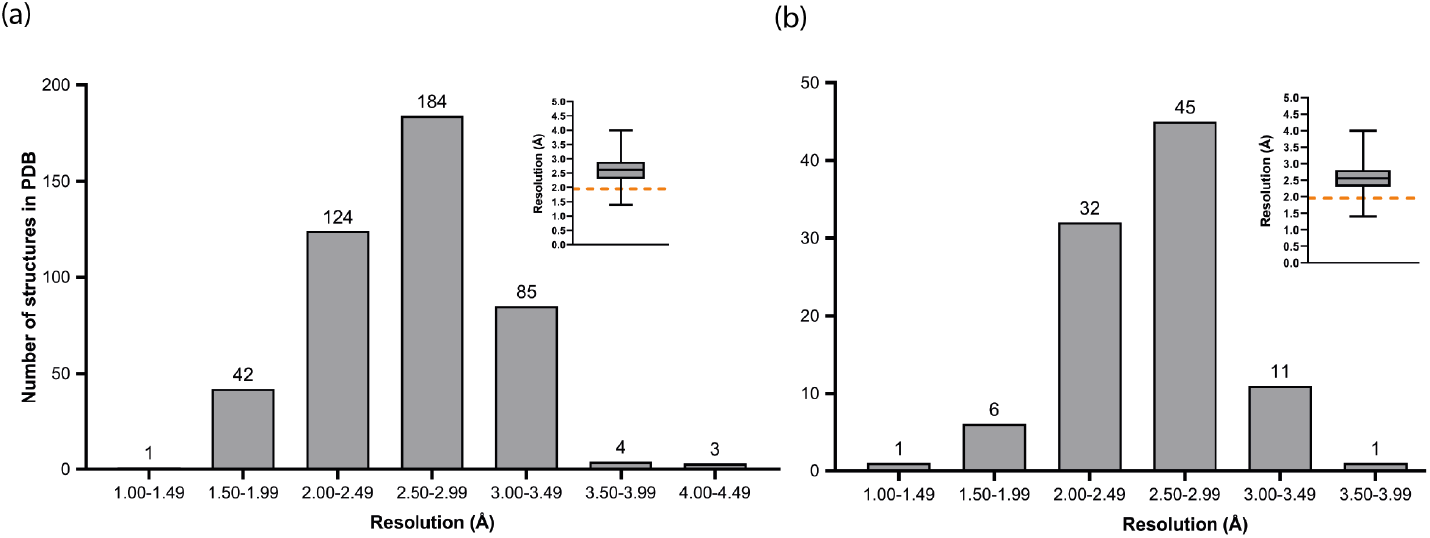
Resolution ranges of deposited TCR:pMHC structures. (a) Distribution of resolutions for X-ray TCR:pMHC structures deposited in the PDB, with the majority of cases (184) resolved at 2.50–2.99 Å, followed by 124 structures at 2.00–2.49 Å. (b) The dataset is limited to TCR:HLA-A*02:01 complexes deposited in the PDB. The highest number of structures (45) is also in the 2.50–2.99 Å bin, followed by 32 in 2.00–2.49 Å. The median resolution is slightly better than in (a), around 2.2 Å, as shown in the box plot inset. Both datasets are dominated by structures with moderate to high resolution (2.0 – 3.0 Å), with very few at extremely high or low resolution. The orange dashed lines in the box plot insets indicate the 1.95 Å resolution of the dataset obtained with the dgTCR-pMHC complex reported here.

Therefore, dgTCRs are promising candidates for structural studies of TCR:pMHC complexes, both as a first choice for the production of soluble TCR, or whenever crystallization trials are harnessed or refolding protocols do not result in sufficient protein yields.

## Discussion

TCR:peptide:MHC interactions are key for T cell antigen recognition, a central process in the adaptive immune system. Therefore, detailed atomic-level characterization of these molecular interactions is a compelling area of investigation for the development of biomedical therapies and biotechnological applications. Indeed, TCRs are a major focus of basic research, particularly for elucidating immune mechanisms, as well as in clinical settings for the treatment of cancer, infections, autoimmune diseases, and transplant rejection.

High-quality TCR samples are sine qua non for high-resolution characterization of TCR structures and TCR:pMHC complex docking and interactions. Although varied eukaryotic expression systems such as insect cells ^17^ and human-derived cell lines ^18^ have been used for the preparation of soluble TCRs for structural studies, refolding of TCR α and β chains from inclusion bodies remains the primary choice. Direct production in bacterial expression systems is limited by the presence of structural intra- and inter-chain disulfide bridges, whose formation is impaired by the reducing environment found in the bacterial cytoplasm. However, viable, soluble TCRs can be obtained by refolding insoluble, unfolded TCR α and β chains from inclusion bodies, as demonstrated by Wiley DC, Garboczi DN and colleagues^12^, and later by Boulter JM, Jakobsen BJ and coworkers^11^. Boulter and Jakobsen’s procedure enabled TCR α/β covalent linkage through artificial cysteines in both chains of the TCR constant regions. All in all, refolding of TCR from inclusion bodies has been replicated by many groups worldwide and still allows a myriad of research studies on the structural biology of the TCR. Nevertheless, refolding of α/β (and γ*/*δ) TCRs consists of multi-step procedures including isolation of denatured inclusion bodies, solubilization, refolding, dialysis, chromatographic and concentration steps to reach the final product. It also involves handling large volumes, particularly during the protein expression and dyalisis stages. Here, considering the amount of protein used for our initial crystallization screenings (0.5 mg), the equivalent of just a 20-ml cell culture was sufficient to produce enough protein for complexation and subsequent crystallization studies.

As for other proteins produced in mammalian cells, TCR chains may undergo several, heterogeneous post-translational modifications (PTMs), such as N-glycosylation. The presence of large, branched sugar molecules protruding from proteins often constrains nucleation, ultimately hindering protein crystallization. For this reason, N-glycosylation removal is a common step applied to mammalian-produced proteins when protein crystallization needs to be achieved. Different strategies can accomplish this task. Enzymatic cleavage is one such option, although it may not remove all the sugar moiety in a homogeneous manner, and additional purification procedures grants counterproductive protein sample loss. Another common option consists of removing the amino acid sequence patterns that drive N-glycosylation, i.e., Asn-X-Thr/Ser (where X is any amino acid). Given their structural analogy, Asn > Gln and Asn > Asp substitution are the most frequent choices for this purpose.

Here, we describe a simple strategy for high-yield recombinant TCR production suitable for crystallization and structural studies. Our strategy prevents N-glycosylation of the TCR alpha and beta constant regions to enhance nucleation and promote crystallization of the target molecules. Human TRAC*01 and TRBC2*01 sequences, commonly used for structural studies, contain three and one N-glycosylation sites, respectively. We have substituted the Asn susceptible sites for Gln residues; more specifically, we performed N28Q, N62Q and N73Q replacements in the TCR alpha chain constant region, and N63Q in the TCR beta chain constant region. This approach enabled the successful production of soluble, non-glycosylated TCR from CHO-S mammalian cells.

Moreover, the deglycosylated TCR strategy resulted in high protein yield, and enhanced crystallization rates of TCR:pMHC complexes. First, dgTCR production in CHO-S cells turned above 26 mg/L of pure protein, whereas production of the wild type equivalent following the same expression procedure rendered 8.4 mg/L. Next, while the deglycosylated TCR:pMHC complex led to 52 initial hits from 384 conditions across four screens, the complex formed with the wild-type TCR produced only one initial hit out of 192 conditions tested (2x 96-well screens). More crystallization conditions should be tested to compare both protein samples and further confirm the observed tendency of the deglycosylated TCR sample to yield higher crystallization rates. In this context, the high crystallization propensity was further supported by new crystallization trials using an additional dgTCR variant with a different CDR3α sequence. Remarkably, this new complex crystallized even more readily than the original, producing crystals in nearly one out of every four drops Additionally, crystals of dgTCR-pMHC were diffraction-ready in as short as 24 hours, as exemplified by the 1.95 Å resolution dataset obtained with a single crystal from the original screen, without further optimizations.

The high-resolution of the obtained dataset enables a precise identification of the peptide amino acids and positioning of the CDR loops at the binding interface, essential for a deep understanding on how pMHC recognition by TCR molecules is driven. In fact, the resolution obtained is higher than the average resolution of other TCR:pMHC complex structures deposited at the PDB, which highlights the utility that our proposed strategy offers for structural analyses of TCR:pMHC interactions.

The high yield and quality of our soluble TCR preparations make them suitable for various structural biology techniques, including cryo-electron microscopy (Cryo-EM) and nuclear magnetic resonance (NMR), in addition to protein crystallography.

All in all, we present a simple and efficient strategy for recombinant TCR production in mammalian cells that avoids refolding and yields high concentrations of monodisperse soluble TCR, promoting high TCR-pMHC complex crystallization rates.

## Methods

### Production and purification of the recombinant TCRs and pMHC

The gene sequences encoding the TCR alpha and beta chains were synthetized by GeneUniversal. The alpha chain was designed to incorporate N146Q, N180Q and N191Q substitutions, a BamHI restriction site between the variable and constant regions, and a C-terminal 6xHis tag. The beta chain construct included an N185Q substitution and a XhoI restriction site located between the variable and constant regions. The sequences were provided in pUC57 vectors flanked by EcoRI and HindIII restriction sites. The pUC57 vectors were digested for 15 minutes at 37 ºC using FastDigest Restriction Enzymes (Fisher Scientific), and the resulting fragments were subsequently cloned in pcDNA3.4 vectors (bioNova) using Optimzyme™ T4 DNA ligase (Thermo Fisher Scientific). Competent *E. coli* DH5α cells were transformed with the ligation products, and positive transformants were grown in 10 mL LB supplemented with ampicillin (50 µg/mL, Fisher). Plasmid DNA was extracted and purified with the GeneJET Plasmid Miniprep Kit (Thermo Fisher Scientific) following the manufacturer’s instructions, and the inserted sequences were verified by Sanger sequencing. To prepare transfection-grade plasmid DNA, the positive DH5α cells were cultured in 100 mL LB under ampicillin (50 µg/mL, Fisher) selection and plasmid DNA was isolated and purified using the PureLink™ HiPure Plasmid Midiprep Kit (Thermo Fisher) according to the manufacturer’s instructions.

CHO-S cell (Invitrogen) were cultured in ExpiCHO™ Expression Medium (Thermo Fisher) and co-transfected with the plasmids encoding the alpha and beta chains using the ExpiFectamine™ CHO Transfection Kit (Thermo Fisher). The transfected cells were then incubated with agitation at 37ºC and 8% CO_2_. Eight days post-transfection, cells were harvested, and the supernatant was dialyzed against 20mM HEPES pH 7.4, 150 mM NaCl (HBS). During the dialysis process, the buffer was exchanged twice daily, using two liters of fresh buffer each time. Prior to purification, the dialyzed sample was filtered using a 0.45 µm filter (Merck) to eliminate any particulate contaminants, and subsequently loaded onto a 1 mL HisTrap affinity chromatography column (Cytiva). The column was washed with 20 mM Hepes pH 7.4, 150 mM NaCl, 20 mM imidazole, followed by elution of the protein with 20 mM Hepes pH 7.4, 150 mM NaCl, 200mM imidazole. Fractions corresponding to the main elution peak were pooled and further purified by size exclusion chromatography on a Superdex 200 10/300 GL column (Cytiva), using TBS (20 mM Tris pH 7.4, 150 mM NaCl) as the mobile phase. TCR-containing fractions were pooled and quantified based on extinction coefficients and absorbance measurements at 280 nm.

The sequences of HLA-A*02:01 and β2 microglobulin were PCR amplified to introduce an N-terminal NcoI restriction site and C-terminal XhoI restriction site. The amplified DNA was digested for 15 minutes at 37 ºC with NcoI and XhoI FasDigest Restriction Enzymes (Fisher Scientific) and subsequently cloned into pET28 vectors using Optizyme™ T4 DNA ligase (Thermo Fisher Scientific). Competent *E. coli* DH5α cells were transformed with the ligation products, and positive colonies were grown in 10 mL LB under kanamycin (50 µg/mL, Panreac AppliChem) selection. Plasmid DNA was extracted and purified using the GeneJET Plasmid Miniprep Kit (Thermo Fisher Scientific) according to the manufacturer’s instructions.

After verifying the sequences by Sanger sequencing, the recombinant plasmids were used to transform *E. coli* BL21(DE3) competent cells (Agilent). Single colonies were cultured in LB with kanamycin and shaken at 37 ºC. When the optical density at 600 nm reached 0.6, protein expression was induced by adding 1 mM isopropyl β-d-1-thiogalactopyranoside (Thermo Fisher Scientific), and the cultures were shaken for an additional 4 hours at 37 ºC.

Bacterial pellets were obtained by centrifugation at 4500 g for 15 minutes and then resuspended in lysis buffer (50 mM Tris pH 8.0, PMSF (Phenylmethanesulfonyl fluoride) 1 mM (Thermo Fisher Scientific), lysozyme 0.25 mg/mL (Sigma), MgCl_2_ 10 mM (Sigma) and DNAase (Sigma). The mixtures were sonicated at 40% amplitude for 5 minutes (30 seconds burst/rest intervals). After sonicating, the samples were centrifuged at 10000 g for 20 minutes, the supernatant was discarded, and the pellets (containing inclusion bodies) were washed once with 50 mM Tris pH 8.8, 10 mM EDTA, and 1% Triton x100 (Sigma) and twice the same buffer without Triton x100. The inclusion bodies were drained of liquid before storage at -20 ºC.

One hundred mg of HLA-A*02:01 and 33 mg of β2 microglobulin inclusion bodies were thawed, solubilized in 8.0 M Urea, 50 mM Tris pH 8.0, 10 mM DTT, and diluted into a 200 mL of refolding buffer (containing 0.4 M L-arginine, 5 mM reduced glutathione and 0.5 mM oxidized glutathione) with 3.6 mg of synthetic peptide (GenScript). The soluble inclusion bodies were refolded by dialyzing against 20 mM Tris pH 8.0 for 72 hours, with the buffer replaced twice daily using two liters of fresh buffer. The dialyzed sample was filtered through a 0.45 µm filter (Merck) to remove particulate matter and then loaded onto a 5 mL HiTrap Q FF anion exchange chromatography column (Cytiva) for purification. Proteins were eluted with a 0-0.4 M NaCl gradient over 20 column volumes. Eluted fractions were assessed by SDS-PAGE and the target (pMHC) samples were pooled and quantified using extinction coefficients and absorbance at 280 nm. All chromatographic procedures were carried out and monitored on an ÄKTA Pure platform (Cytiva).

The ternary TCR:pMHC complexes were prepared by mixing stoichiometric amounts of either wild type or dgTCR and pMHC. The final protein samples were concentrated to 5 mg/mL using a Nanosep 30 kDa cutoff device (VWR) for crystallization trials.

### Crystallization of the TCR:pMHC complexes

The ternary TCR:pMHC complex was screened against using the sitting drop vapor diffusion method and 20 ºC incubation. A Crystal Gryphon LCP platform (Dunn Labortechnik GmbH) was used for these trials. Drop volumes were set up to 300 nanoliters for the crystallization reagent in each condition, while the protein sample drop volumes ranged 250 to 300 nanoliters. Initial hits were optimized to obtain larger crystals, whichwere harvested, soaked in crystallization medium supplemented with 20% glycerol or 20% ethylene glycol, and cryo-cooled in liquid nitrogen before diffraction analyses.

### Diffraction analysis, data processing and structure determination and refinement

X-ray diffraction and data collection were performed on the BL13-XALOC beamline at the Alba synchrotron facility (Cerdanyola del Vallès, Barcelona, Spain). The best crystals, which allowed collection of full datasets at 1.95 Å were grown in 0.2 M NaCl, 0.1 M MES pH 6.0 and 20% PEG 6000. The diffraction data was reduced and integrated with XDS ^19^ and then merged and scaled with *AIMLESS* ^20^ in the CCP4 Suite ^21^. A 5% of reflections were excluded for validation purposes during the refinement process. The structure was solved by molecular replacement *via* Phaser ^22^. Five independent templates were used for phasing. First, the MHC class I molecules of HLA-A*02:01 and β2 microglobulin from PDB IDs 7KGP and 7EJL, respectively, the human constant domains from PDB ID 7Z50, and lastly the variable domains of alpha and beta chains from PDBs 3TJH and 8RO5, respectively. Molecular replacement solutions were subsequently refined with Refmac5 ^23^ or phenix.refine ^24,25^. The final molecule was generated after iterative cycles of manual building in *Coot* ^26^ and further refinement.

## Acknowledgements

We are grateful to the staff of the Xaloc beamline at ALBA Synchrotron for their assistance with X-ray diffraction data collection. This work was funded by Gobierno de Navarra Ayudas para la realización de Proyectos Estratégicos de I+D 2023-2026 (0011-1411-2023-000072, PITAGORAS) and Ayudas a los agentes del SINAI para realizar proyectos de I+D colaborativos 2024 (Grant PC24-MEMORIAL-017-003-015).

## Author contributions

Conceived research: JLS; Performed experiments: LOO, EEA, MGDR, JLS, SH; Data analysis: JLS, EEA, LOO; Draft writing: EEA, JLS, LOO.

